# Optimal strategies to protect a sub-population at risk due to an established epidemic

**DOI:** 10.1101/2021.09.10.459742

**Authors:** Elliott H. Bussell, Nik J. Cunniffe

## Abstract

Epidemics can particularly threaten certain sub-populations. For example, for SARS-CoV-2, the elderly are often preferentially protected. For diseases of plants and animals, certain sub-populations can drive mitigation because they are intrinsically more valuable for ecological, economic, socio-cultural or political reasons. Here we use optimal control theory to identify strategies to optimally protect a “high value” sub-population when there is a limited budget and epidemiological uncertainty. We use protection of the Redwood National Park in California in the face of the large ongoing state-wide epidemic of sudden oak death (caused by *Phytophthora ramorum*) as a case study. We concentrate on whether control should be focused entirely within the National Park itself, or whether treatment of the growing epidemic in the surrounding “buffer region” can instead be more profitable. We find that, depending on rates of infection and the size of the ongoing epidemic, focusing control on the high value region is often optimal. However, priority should sometimes switch from the buffer region to the high value region only as the local outbreak grows. We characterise how the timing of any switch depends on epidemiological and logistic parameters, and test robustness to systematic misspecification of these factors due to imperfect prior knowledge.

## Introduction

Management of emerging infectious disease is most likely to be successful when it starts as soon as possible. Smaller epidemics are easier and less expensive to control than the larger epidemics which would result if management were to be delayed (***Althaus et al., 2015***; ***Epanchin-Niell and Hastings, 2010a***; ***Hoffman and Silverberg, 2018***; ***Longini et al., 2015***). Sufficiently rapid intervention can even make eradication, or at least localised elimination, a realistic proposition (***Dowdle, 1998***; ***Klepac et al., 2015***). Our recent experience with COVID-19 provides an object lesson. Governments of some countries in the Asia-Pacific region took early and decisive action to introduce non-pharmaceutical interventions (***Pearce et al., 2020***; ***Thompson et al., 2020b***). Certain of these countries, perhaps most notably New Zealand, therefore appear to continue to be in a very good position to face the ongoing challenge of the global pandemic (***Summers et al., 2020***). In particular, despite recent incursions (***Dyer, 2021***), they seem better-placed than any other nation to enact the “Zero COVID” strategy based on prompt responses to any incursion, with no tolerance for community transmission (***Baker et al., 2020***).

However rapid responses are not always possible. Pathogens can become established in a host population without the diseases they cause being identified, particularly when effective surveillance systems are not in place (***Carvajal-Yepes et al., 2019***; ***Grogan et al., 2014***; ***Mastin et al., 2020***; ***Ristaino et al., 2021***) or if there is a long incubation period before symptoms (***Thompson et al., 2016b***; ***Khan et al., 2020***). Attempts to eliminate or eradicate are also not always successful (***Thompson et al., 2020a***). If transmission is possible before – or perhaps even without – infected hosts showing symptoms, disease management is difficult (***Cunniffe et al., 2015b***; ***Cunniffe and Gilligan, 2020***; ***Fraser et al., 2004***). Early in any outbreak it might also be unclear which controls or combinations of controls are likely to be most successful (***Thompson et al., 2018***). Particularly for plant and animal pathogens, there are often also economic constraints which mean that extensive and costly management is simply not justified (***Wilkinson et al., 2011***).

Strategies to effectively mitigate well-established outbreaks are therefore very important. Plant disease epidemics provide a pressing example. The impacts of plant pathogens on food security (***Strange and Scott, 2005***) and ecosystem services (***Boyd et al., 2013***) are well acknowledged. However, management of plant disease epidemics is very often a case of “too little too late” (***Tomlinson and Potter, 2010***). For tree diseases, delays and/or deficiencies in detection have been implicated in high-profile failures of various large-scale control programs, including for chestnut blight (***Freinkel, 1997***), white pine blister rust (***Maloy, 1997***), Dutch elm disease (***Tomlinson and Potter, 2010***) and citrus canker (***Gottwald, 2007***). Indeed, elimination is often not even attempted. For example, following the first detection of ash dieback (a disease of ash trees caused by the fungus *Hymenoscyphus fraxineus*) into the United Kingdom in late 2012, a consensus was rapidly formed that country-wide management was unlikely to succeed because the pathogen was already so widely-dispersed (***Thomas, 2016***). In Italy, control of olive quick decline syndrome (caused by the bacterial pathogen *Xylella fastidiosa*) is based entirely on slowing the spread, with management and detection focused on “buffer” and “containment” zones bordering an “infected” zone within which the disease is, essentially, allowed to spread undetected and uncontrolled (***EFSA Panel on Plant Health (PLH) et al., 2019***).

We take control of sudden oak death in coastal forests of central California to southwestern Oregon, caused by the oomycete *Phytophthora ramorum*, as an example of a well-established plant disease epidemic that still requires management. The generalist pathogen *P. ramorum* affects over one hundred plant species, and broadly-speaking causes two kinds of disease in the area in question: “sudden oak death” on tanoaks and oaks (including coast live oak), and “ramorum blight” on a large number of species of woody shrubs and forest understorey plants (***Grünwald et al., 2019***). Sudden oak death causes large bleeding cankers to form on the main stem of affected trees, eventually leading to death (***Rizzo et al., 2005***). However, some species affected by ramorum blight, in California most notably bay laurel, are not killed by the infection. They therefore act as “spreader species” by supporting significant foliar sporulation over extended periods (***Grünwald et al., 2008***). The pathogen spreads predominantly through short-distance rain splash dispersal of spores, but spores can be dispersed over longer distances by turbulent air currents, rivers and streams, or when carried by animals or human activity (***Grünwald et al., 2012***).

A very large sudden oak death epidemic has devastated coastal forests of central California to southwestern Oregon since its first detection in California in 1995, killing millions of oak (*Quercus* spp.) and tanoak (*Notholithocarpus densiflorus*) trees (***Meentemeyer et al., 2011***). That epidemic is now very widespread, covering over 2000km^2^ in California alone (***Grünwald et al., 2019***; ***Peterson et al., 2015***) (Figure 1a). Economic impacts are significant, with, for example, an estimated $135M USD loss in property values attributed to the disease (***Kovacs et al., 2011***). Recent large-scale modelling work has shown that successful control of sudden oak death in California is no longer possible, and indeed has been impossible for many years (***Cunniffe et al., 2016***).

**Figure 1.**
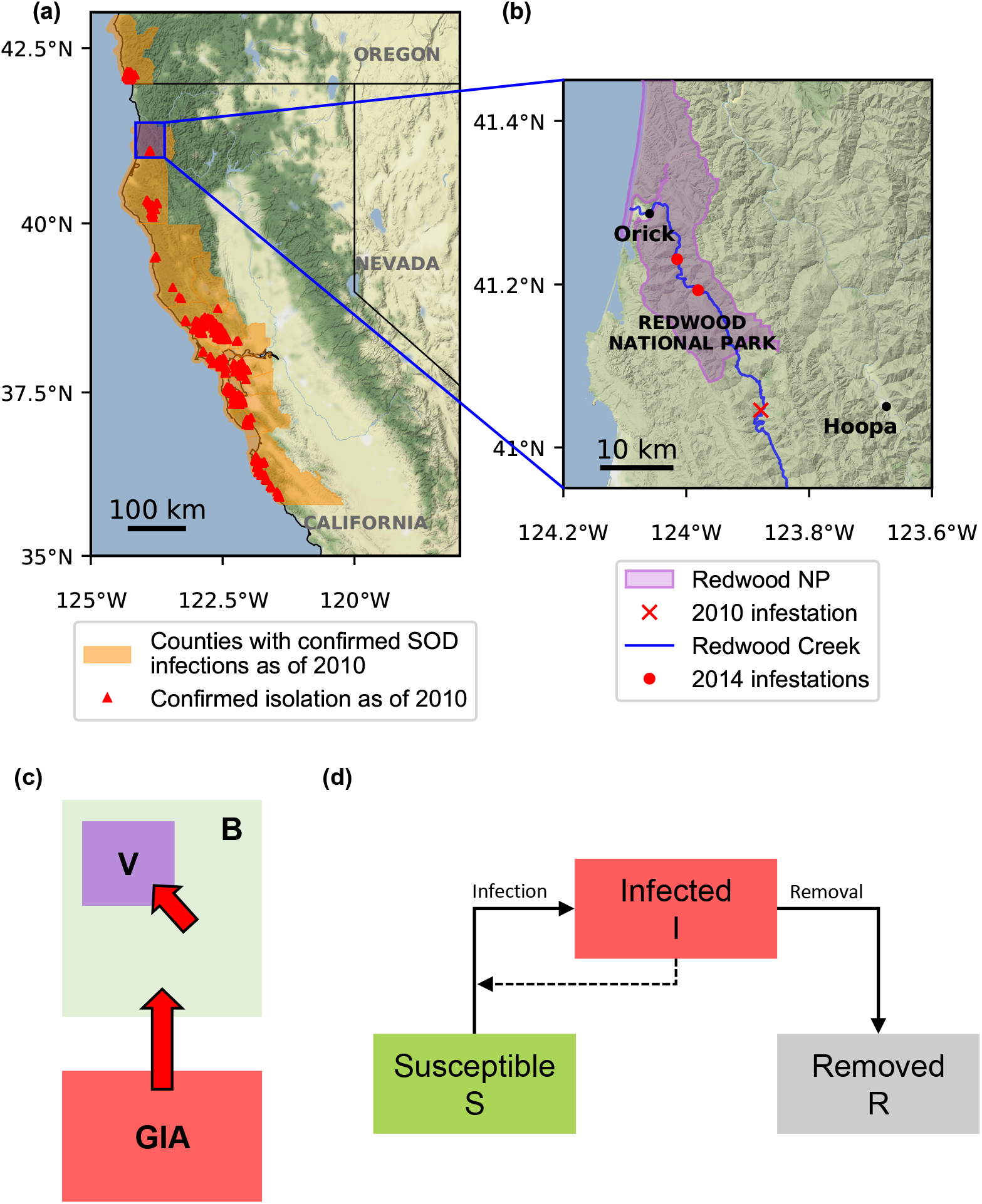
Maps showing areas with confirmed sudden oak death infections, and schematics of our model’s spatial structure and within-patch dynamics. **(a)** Confirmed sudden oak death infections in California in 2010 (infected counties as specified in ***Grünwald et al.*** (***2019***), with locations of confirmed isolations taken from the SODmap project (***Garbeletto et al., 2014***)). The bulk of the epidemic lies in a relatively contiguous and heavily infested area of coastal California centred roughly on latitude around 37.5°N, near to the sites of first detection of the disease in Marin and Santa Cruz counties. More northerly outbreaks have thus far remained more spatially distinct. **(b)** Area near to Redwood National Park, showing the location of the initial incursion as well as the later detections of symptoms within the borders of the National Park itself (locations of infestations in 2010 and 2014 again taken from ***Garbeletto et al. (2014)***). **(c)** The spatial structure of the system as modelled. A generally infested area (GIA, which here corresponds to the rest of California) provides a source of inoculum, generating a constant force of infection on a buffer region which is only very lightly infected (B, which here corresponds to Redwood Creek). The infection can then in turn invade the high value region (V, which here corresponds to the Redwood Creek National Park). The goal of control is to protect the hosts in the high value region. **(d)** Within each region the epidemic follows SIR dynamics, where hosts move from Susceptible (S) to Infected (I) and are Removed by the disease or through control (R). The dashed line indicates that the number of infected hosts influences the infection rate; removal includes components corresponding to both disease-induced host mortality as well as roguing (i.e. removal of infectious hosts as a form of disease control).

However, control of isolated outbreaks and localised management both remain firmly on the agenda. In 2001 a spatially-distinct epidemic was identified in Curry County, Oregon, and since then over $20 million USD has been spent on identifying and treating that localised epidemic (***Grünwald et al., 2019***). There are also other areas ahead of the main epidemic front under active management, most notably a pair of relatively large outbreaks in Humboldt County, in California (Figure 1a; the southern border of Humboldt County is at a latitude of around 40°N). Although further south than the outbreak in Oregon, the outbreaks in Humboldt county are some distance “ahead” of the main bulk of the epidemic (***Alexander and Lee, 2010***).

Disease management in Humboldt County exemplifies the challenges now posed by the control of sudden oak death. The goal is to design a management scheme that can effectively achieve a smaller, more local, objective than complete elimination or eradication. This could be, for example, slowing local rates of disease spread or protection of valuable resources (***Cobb et al., 2013***). This must be done when there is a limited budget available for control (***Cunniffe et al., 2016***). We focus here on strategies which reduce impacts of disease on a particular “high value” region, within which it is important to mitigate the effects of disease for ecological, economic, socio-cultural or political reasons.

For sudden oak death in California, any high value region would most obviously be defined in terms of an area with particular ecological and/or socio-cultural value. The host species that is most affected – tanoak – is not grown for its timber. Disease impacts therefore predominantly include deleterious effects upon carbon sequestration, fuel dynamics and cultural resources. However, direct economic effects due to sudden oak death have been documented in California, most notably effects on property values (***Kovacs et al., 2011***). Effects on timber production are also relevant in other areas. For example, emerging *P. ramorum* lineages in Oregon might also potentially threaten Douglas-fir silviculture. The epidemic caused by *P. ramorum* in the United Kingdom is most often known as “sudden larch death”, since the coniferous plantation species Japanese larch *Larix kaempferi* is predominantly affected in that context.

Although other controls such as chemical treatments have occasionally been promoted for *P. ramorum* (***Alexander and Lee, 2010***), and despite targeted prophylactic removal of certain host species sometimes being recommended as a potentially highly effective strategy to reduce risks of sudden oak death epidemics (***Swiecki and Bernhardt, 2013***), there is a consensus that effective disease management at medium to large spatial scales must be based upon removal of infected hosts (***Grünwald et al., 2019***; ***Cunniffe et al., 2016***). The only outbreak of sudden oak death that is actively being controlled across large spatial scales in the United States, in Curry County, Oregon, is certainly being managed in this way (***Peterson et al., 2015***). We therefore concentrate exclusively on this type of control here.

The particular example we select to motivate our analysis is based upon the Redwood Creek outbreak (Figure 1b). This outbreak was detected in May 2010 through stream monitoring, with the disease found to be present in a stream near Orick, at the mouth of Redwood Creek (***Valachovic et al., 2013***). It was considered to be an outbreak of high significance due to its proximity to Redwood National Park as well as the traditional and reservation lands of the Yurok and Hoopa tribes, respectively. Extensive detection efforts were carried out to identify the source of the infection, which was located in Redwood Valley in July 2010 (***Stark et al., 2014***). Disease was also detected within the National Park within a few years of the initial incursion into Redwood Creek (Figure 1b). Treatment aiming to prevent further damage to Redwood National Park is ongoing (***Croucher et al., 2013***). A pressing question is how limited control resources should be partitioned to protect the high value region, i.e. the Redwood National Park. The most obvious, and pressing, question is whether management resources should be entirely focused on treating within the high value region itself, or whether treatment of the growing epidemic in the surrounding areas can better serve the goal of protecting the National Park.

We show in principle how control strategies to protect the high-value region can be designed and tested using a mathematical model. The role of modelling in optimising decision making in disease control is now well-established (***Lofgren et al., 2014***), including in plant health (***Cunniffe and Gilligan, 2020***). Models offer a rational basis to decide where, when and how to control disease outbreaks (***Cunniffe et al., 2015a***). However, the conventional approach for testing interventions via models - comparing a relatively small number of pre-specified intervention strategies - necessarily risks sub-optimal performance, since only a small subset of all possible strategies can ever be tested in depth (***Cunniffe et al., 2016***). Recent work has shown how the search space of possible interventions can be unambiguously explored for plant diseases using optimal control theory (***Bokil et al., 2019***; ***Bussell et al., 2019***; ***Bussell and Cunniffe, 2020***; ***Dangerfield et al., 2019***; ***Hamelin et al., 2021***; ***Ndeffo Mbah and Gilligan, 2010b***,a, ****2011****).

We concentrate here upon how optimal control theory can be used to solve the problem of prioritising different areas for disease management. Our analysis is a proof of concept, capturing the key features of the Redwood Creek outbreak of sudden oak death in a simple way. Indeed, while detailed models of this system are possible (***Cunniffe et al., 2016***; ***Bussell and Cunniffe, 2020***), here we instead concentrate on a rather general model, which in principle is applicable to many host-pathogen combinations. We test the performance of optimal control theory in optimising management in this model, performing extensive scans over different values of epidemiological and logistical parameters, ensuring a wide range of scenarios are captured. Since we deliberately phrase our mathematical model to omit many system specific details, our results therefore illustrate epidemiological principles important whenever a subset of host individuals must be protected in the face of a large, ongoing epidemic.

In particular we use optimal control theory to understand how time-dependent disease management can be optimised in order to protect a high value region at risk from a growing, spreading epidemic. We account for economic and logistical limitations in control by introducing a budgetary constraint such that only a finite number of infected hosts can be treated per unit time (***Ndeffo Mbah and Gilligan, 2011***; ***Dangerfield et al., 2019***). We then seek to understand how these finite resources for disease management should be partitioned, and whether and how the balance of control effort within *versus* outside the high value region changes over time and with the state of the epidemic. Management strategies as derived using optimal control theory can often be rather complex (***Bussell et al., 2019***; ***Bussell and Cunniffe, 2020***). We therefore focus here on how optimal strategies can be characterised in an intuitive and understandable fashion. This would be a precondition of implementation by local managers and other stakeholders. We also use our model to understand how optimal management strategies would be affected by alterations to, or imprecision in our knowledge of, epidemiological and/or logistic parameters, commonly the case in tackling an emerging epidemic (***Thompson et al., 2018***).

## Methods

### Epidemiological model

We split the host landscape into three disjoint regions: a generally infested area in which the disease is already well-established, a buffer region in which the disease might perhaps be present, but has not yet become established, and a high value region that is currently entirely uninfected and that must be protected. In the context of Redwood Creek and sudden oak death, the generally infested area would correspond to the large epidemic across large parts of California (***Alexander and Lee, 2010***), the buffer region would be the recently-infected forests of the Redwood Creek watershed (***Stark et al., 2014***), and the high value region would be the Redwood National Park (***Croucher et al., 2013***) (Figure 1b,c). To reduce the number of state variables and parameters in our model, we simplify the epidemic in the generally infested area such that it is represented as a source of external inoculum, generating a constant force of infection upon the buffer region (in general the “force of infection” is the per capita rate at which susceptible hosts become infected (***Keeling and Rohani, 2008***); here we assume the component of this corresponding to the epidemic elsewhere remains constant over time). This simplification is valid when considering new, relatively isolated outbreaks of disease: a common situation for diseases such as sudden oak death that are spread over long distances through rare long-distance dispersal events (***Meentemeyer et al., 2011***). As shown in electronic supplementary material, S1 text, the effects of even a growing rather than constant-sized external epidemic upon the dynamics within the region of interest can also often be subsumed into an appropriately time-averaged external force of infection. Since we scan over a range of values of the constant force of infection in the results we present, we consider the simplification we make to use only constant values of this parameter to therefore not be unduly restrictive.

The buffer and high value regions are modelled as well-mixed patches, meaning the only spatial component in our model is between-patch coupling. We use a (*S*)usceptible-(*I*)nfected-(*R*)emoved model for the epidemic dynamics in each patch (***Keeling and Rohani, 2008***), in which hosts can be susceptible to the disease (*S*), infected and infectious (*I*), or removed (*R*) (Figure 1d). We assume that infection can potentially kill hosts, meaning an infected host can be removed either by disease-induced death or via disease control. Guided by the epidemiology of sudden oak death, for which standing dead trees are not considered to be a significant source of inoculum (particularly in comparison to foliar hosts which are not killed by infection) we assume that removed hosts cannot cause other hosts to become infected (***Rizzo et al., 2005***). We seek to optimise management of the epidemic by determining time-dependent allocations of control resource to the buffer and high value regions, treating a time-varying proportion of the infected hosts in each region. This control, which for plant pathogens is often known as “roguing” (***Cunniffe et al., 2014***), is therefore the only form of disease management we allow for in our model.

Each patch has a particular fixed population size (*N*_*B*_ and *N*_*V*_ for the buffer and high value regions, respectively), and the two patches are linked for transmission, symmetrically, by a coupling constant *ϵ*. Taking the within-patch transmission rate to be *β*, and assuming density dependent transmission (***Keeling and Rohani, 2008***), we obtain the following system:

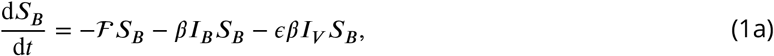

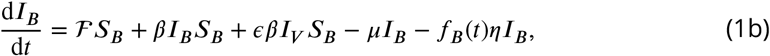

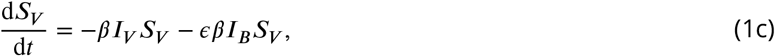

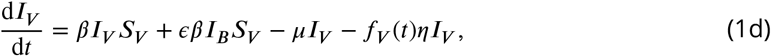

in which the subscripts *B* and *V* refer to the buffer and high value regions, respectively. The (constant) external force of infection from the generally infested area is given by 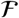, and we assume only the buffer region can be thus infected. The infectious period in the absence of control is assumed to be 1/*μ*. Infected hosts can also be removed by roguing at an additional rate *η*. The control inputs *f*_*B*_(*t*) and *f*_*V*_(*t*) are the time-dependent proportions of infected hosts that are being controlled at time *t* in the buffer and high value regions, respectively. It is the functions *f*_*B*_(*t*) and *f*_*V*_(*t*) that we seek to identify in our optimal control problem.

### Optimal control problem

The objective of control is to limit the pathogen’s impacts on the high value region. In particular, we aim to minimise the number of infected and removed hosts in the high value region at some terminal time *T*. Since our model does not include host demography, this is equivalent to maximising the number of susceptible hosts retained in the high value region at this time.

We capture the economic and logistical limitations of disease management by restricting the total number of hosts that can be rogued per unit time (over both regions). The maximum expenditure rate, i.e. the largest number of hosts that can instantaneously be rogued, is assumed to be *M*. This gives the following optimal control problem:

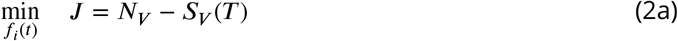

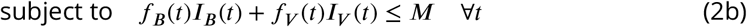

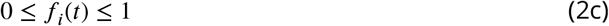

in which the state dynamics are controlled by Equations 1 and the index *i* ∈ {***B, V***}.

We solve the optimal control problem using a direct formulation (***Betts, 2010***). In particular, we time-discretise values of the state and control variables, and treat these as optimisation variables in a nonlinear programming (NLP) problem. State dynamics and initial conditions are included as constraints on the NLP variables, and the optimisation is carried out to minimise the objective value.

We use the BOCOP package (v.2.0.5) (***Team Commands, Inria Saclay, 2017***) to generate the NLP problem, coding the state dynamics (Equations 1) in C++. The package automates discretisation of the system using a fourth order Runge-Kutta method. The NLP problem that results is solved using the software Ipopt (***Wächter and Biegler, 2006***), which implements an interior point optimisation method. The BOCOP software allows extraction of the Lagrange multipliers associated with the state dynamics, i.e. the co-state or adjoint dynamics of the optimal control problem. Using these the relative importance of control in each region can be derived since the higher the co-state, the more benefit to the objective there is from control in that region. This gives the control priority of each region over time, which characterises the overall control policy in readily understandable terms.

### Model parameterisation

Although our modelling was motivated by the spread of sudden oak death in the vicinity of Redwood National Park, our focus here is not to develop detailed control recommendations for any particular application. Our numerical work instead concentrates upon characterising the broad features of optimal controls, and how these features are conditioned on parameter values. We therefore identify a plausible, although arbitrary, parameterisation of our model (Table 1), and use it to drive our analysis.

**Table 1.** Definitions of parameter and state variables, with biological meanings and default values.

| Symbol | Meaning | Default | Sensitivity scans |
| --- | --- | --- | --- |
| $N_B$ | Number of hosts (buffer region) | 500 hosts | |
| $N_V$ | Number of hosts (high value region) | 100 hosts | |
| $S_B$ | Number of susceptible hosts (buffer region) | n/a | |
| $S_V$ | Number of susceptible hosts (high value region) | n/a | |
| $I_B$ | Number of infected hosts (buffer region) | n/a | |
| $I_V$ | Number of infected hosts (high value region) | n/a | |
| $f_B(t)$ | Proportion rogued at time $t$ (buffer region) | n/a | |
| $f_V(t)$ | Proportion rogued at time $t$ (high value region) | n/a | |
| $J$ | Objective function | Equation 2a | |
| $M$ | Maximum expenditure rate | 10 hosts | Figure 4a |
| $\beta$ | Infection rate | $0.005 \text{ host}^{-1} \text{ t}^{-1}$ | Figures 3, 4 and 5 |
| $\eta$ | Roguing rate | $0.2 \text{ t}^{-1}$ | Figure 4b |
| $I_B(0)$ | Initial number of infected hosts (buffer region) | 5 hosts | Figure 5a |
| $\mathcal{F}$ | External force of infection | $0.0 \text{ t}^{-1}$ | Figure 5b |
| $I_V(0)$ | Initial number of infected hosts (high value region) | 0 hosts | |
| $T$ | Time horizon of interest | 12.5 t | |
| $\epsilon$ | Coupling between regions | 0.3 | |
| $\mu$ | Pathogen-induced death rate | $1.0 \text{ t}^{-1}$ | |

We follow previous modelling work targetting the sudden oak death system (***Meentemeyer et al., 2011***; ***Thompson et al., 2016a***; ***Cunniffe et al., 2016***) in using numbers of “hosts” as a convenient shorthand for the appropriate modelling quantum, which almost always corresponds to a relatively large area of contiguous vegetation, and which accounts implicitly for species-specific differences in infectivity and susceptibility. The unit used to measure host abundance is scaled into the numerical values of the infection rate (*β*) and maximum budget (*M*), anyway, and so the dimensions of these quantities do not affect our results. We scale time by the infectious period in the absence of control, allowing us to fix the disease-induced removal rate to be *μ* = 1 in all numerical work.

### Scenarios tested

In the absence of any budgetary constraint, it might be expected that control would be maximal at all times throughout the epidemic in both regions (i.e. *f*_*B*_(*t*) = *f*_*V*_(*t*) = 1 for all *t*). We verified this was indeed the case in our exploratory work by running our optimisation procedure using our default parameterisation, but with the maximum budget, *M*, set to be a very large value. We therefore reverted to a more limited budget for our remaining numerical work, and first concentrated on identifying broad classes of control strategy. As we describe below, these could conveniently be characterised in terms of which region was prioritised for control at which points of the epidemic. We then considered how the details of optimal management strategies were affected by the values of epidemiological and logistic parameters. Finally we investigated the robustness of our results, by examining the performance of optimal controls as derived from a model in which the infection rate was systematically misspecified.

## Results

### Optimal strategies can be characterised in terms of which region(s) are prioritised for control

#### Prioritising the high value region for all time

For our default parameter set (Table 1), the optimal strategy prioritises control of infection in the high value region throughout the entire epidemic (Figure 2). However, since the cost of control within the high value region at any given time depends on the number of infected hosts it then contains, often the high value region can be treated at the maximal rate(i.e. *f*_*V*_(*t*) = 1), but some budget nevertheless remains unspent(i.e. *f*_*V*_(*t*)*I*_*V*_(*t*) = *I*_*V*_(*t*) *< M*). During these periods the optimal strategy diverts any remaining resources to the buffer region(i.e. *f*_*B*_(*t*) *>* 0 for these times).

**Figure 2.**
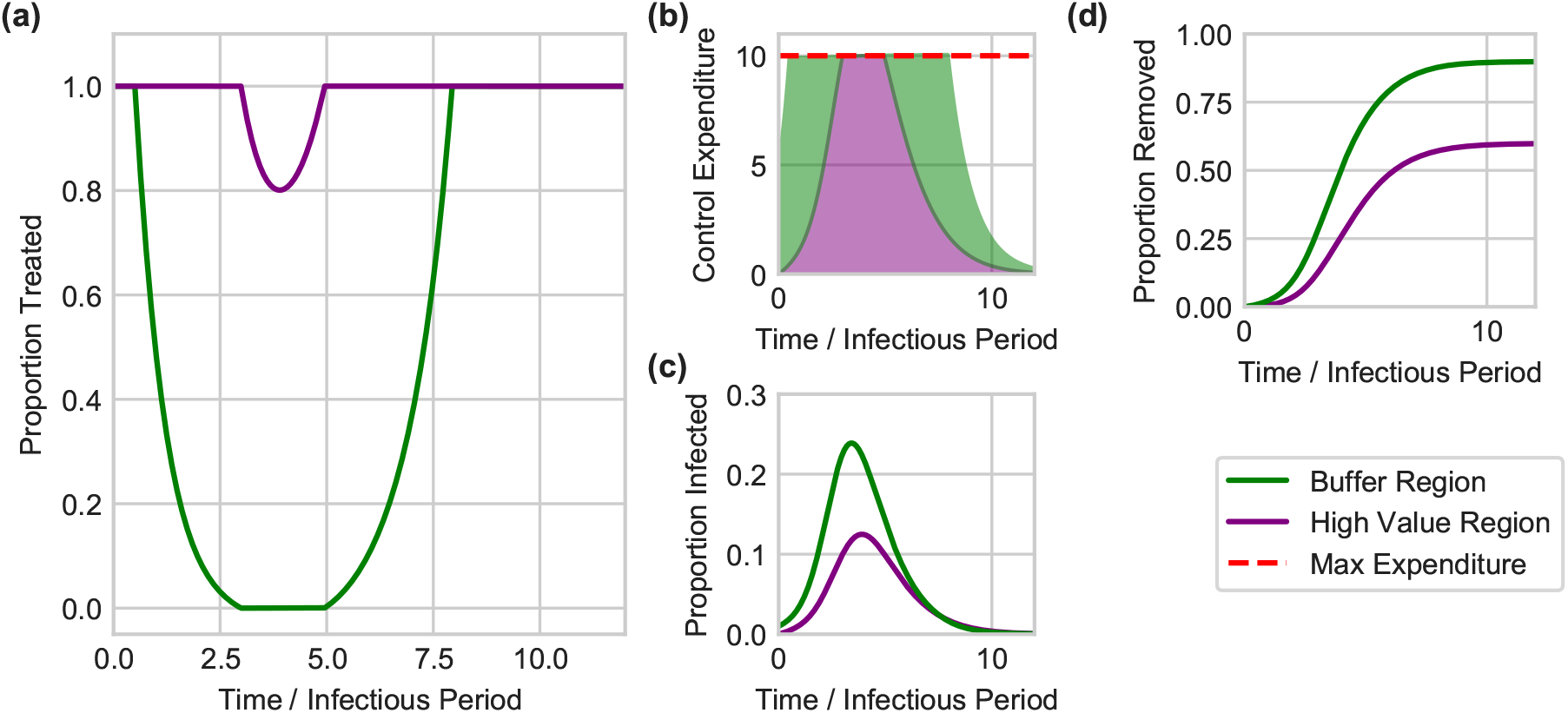
Optimal control can sometimes focus disease management upon the high value region throughout the entire epidemic. The optimal strategy as found for the default model parameterisation(Table 1). **(a)** The proportion of hosts treated in each region over time. **(b)** The number of hosts being treated – which corresponds to the expenditure – in each region over time. The optimal strategy allocates as many resources as possible to the high value region, before spending any remainder in round 3 and 5 time units the cost of maximally removing infected hosts in the high value region would exceed the maximum permissible expenditure, *M*, and so a proportion of infected hosts go untreated even in the high value region. (c) The time-dependence in the proportion of hosts infected in both regions. **(d)** Proportions of hosts that are removed (either by disease-induced death or by control).

However, for our default model parameterisation, there is also a period in the vicinity of the epidemic peak during which there is insufficient resource to fully control all infected hosts in the high value region(i.e. *I*_*V*_(*t*) *> M*). Infected hosts within the high value region then cannot be treated at the maximum rate(i.e. *f*_*V*_(*t*) *<* 1), since it would be too expensive. For the default parameterisation of our model, this occurs at around 3 *< t <* 5 time units.

In general, for strategies which focus upon the high value region throughout the epidemic, whether or not there is such a period within which the budget would be exceeded by fully controlling the epidemic in the high value region depends on the interplay between the maximum budget and the dynamics of disease(see below).

#### Switching focus from the buffer to the high value region

For other parameter sets, however, the optimal strategy switches focus between regions as the epidemic progresses(Figure 3 shows results from a model parameterisation for which this occurs). Early in the epidemic it is optimal to prioritise the buffer region, since doing so slows spread into the high value region. However, later in the epidemic the high value region is again prioritised(although for these parameters there is still some budget to partially treat the buffer region, even when the high value region is treated as aggressively as possible, i.e. *f*_*V*_(*t*) = 1).

**Figure 3.**
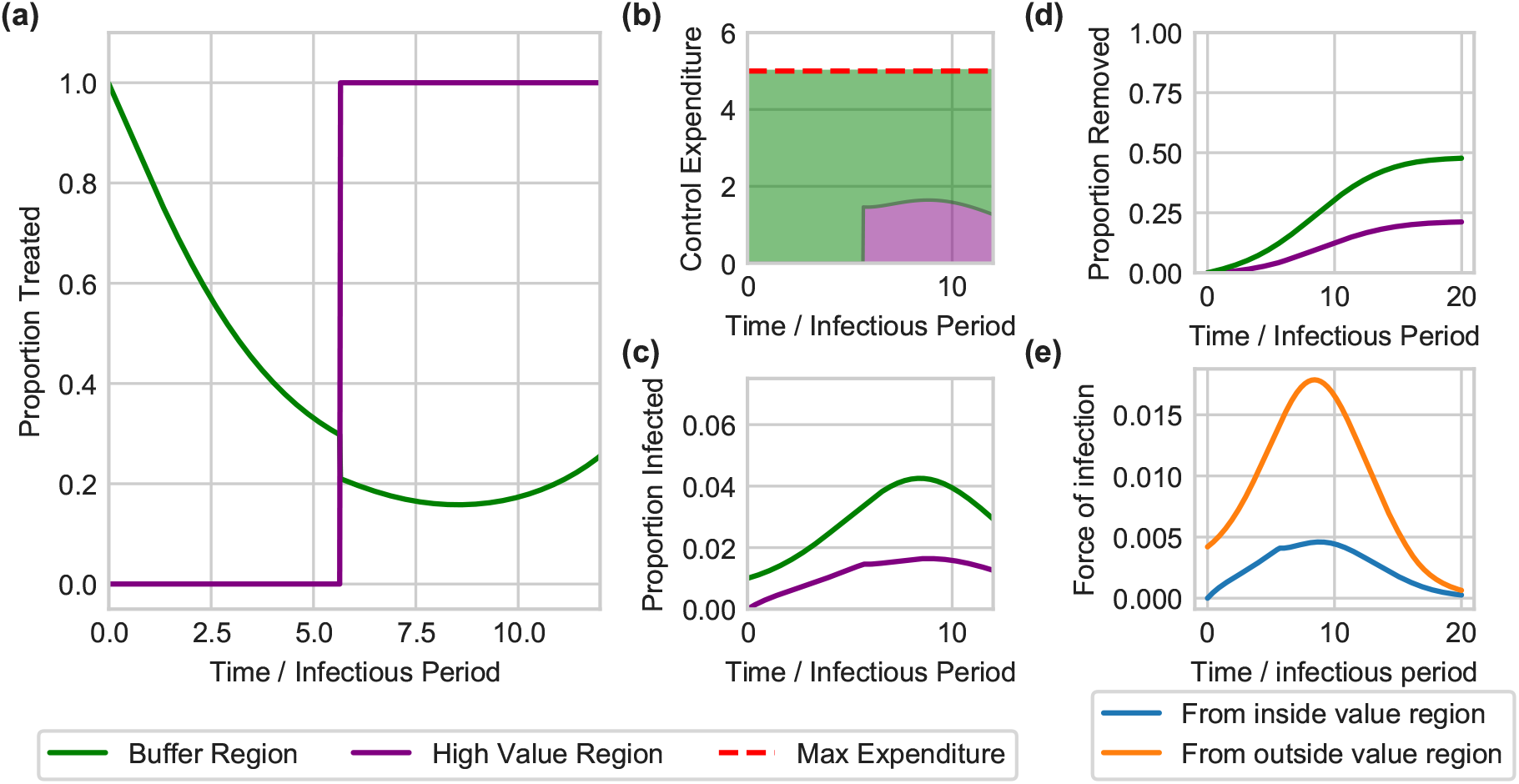
An alternative optimal strategy initially focuses management upon the buffer region, but switches to concentrate roguing within the high value region as the epidemic takes hold. The optimal strategy as found for a model parameterisation slightly altered from the default in Table 1, with a smaller infection rate(*β* = 0.0028 host^−1^ t^−1^) and smaller maximum budget (*M* = 5 hosts). **(a)** and **(b)** show the proportion and number treated over time in both regions. For these parameters, for approximately the first 6 units of time, the buffer region is prioritised, before the optimal control strategy switches to prioritising the high value region thereafter. Disease progress curves are shown in **(c)** and **(d)**. **(e)** shows the force of infection in the high value region from inside and outside the same region. Since the largest part of the infectious pressure on the high value region comes from the buffer region throughout the entire epidemic, the identity of the region providing the dominant infectious pressure cannot be the sole driver of the switch.

A naïve expectation might be that the switch is driven by the optimal strategy targeting whichever region generates the larger force of infection on the high value region at any time. However, the switch time is not simply when the force of infection upon the high value region from within itself becomes greater than that from outside. This is exemplified by Figure 3(e), which shows that even for this parameterisation, the force of infection upon the high value region from itself remains smaller than that from the buffer region throughout the epidemic, despite the switch in focus of the optimal control at *t* ≈ 6 units of time.

Further switches, i.e. multiple changes in focus between the two regions, can occur whilst budgets are not limiting. In these cases since the budget is not limiting, control can be maximal in both regions and so the switch has no effect on the control performed. For the range of parameters we considered here, we do not see strategies with multiple switches which impact the control performed.

### The optimal strategy depends on epidemiological and logistic parameters

#### Switching focus between regions is often optimal at “intermediate” transmission rates

The switching strategy exemplified by Figure 3 is promoted by intermediate values of the infection rate, *β*, although the precise meaning of “intermediate” is heavily conditioned upon the maximum budget, *M* (Figure 4a).

**Figure 4.**
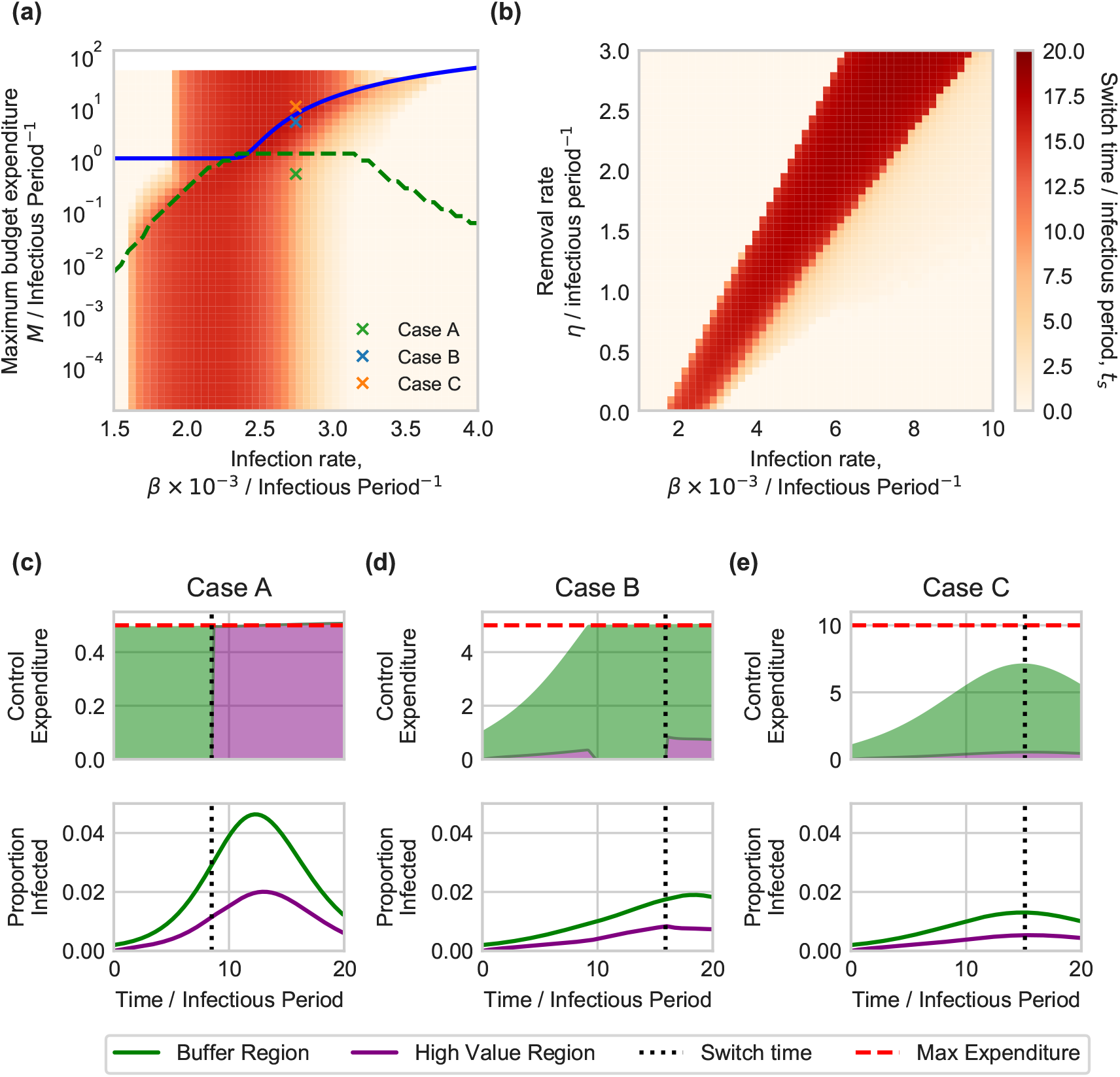
Effect of the infection rate, maximum budget and rate of control on the optimal switching time. Parameter scans were run relative to a baseline set by the default parameters in Table 1, although with only a single initially infected host in the buffer region (*I*_*B*_(0) = 1) and using a longer timescale of interest(*T* = 20) to allow patterns in switching times to be more clearly seen. **(a)** and **(b)** show the effect of infection rate(*β*) combined with the maximum budget(*M*) and removal rate(*η*) respectively. In all cases switching is optimal for intermediate values of the infection rate, *β*, although as control becomes more effective the range of infection rates over which a switch is optimal is shifted to higher values. The “kink” in(a) is caused by whether or not the maximum budget is exceeded by fully treating the high value region(see main text). The blue line indicates where the budget becomes limiting on control resources, i.e. below the blue line maximal control can be carried out in both regions for the full epidemic. The green dashed line indicates where the budget becomes limiting for the full simulation time, i.e. below this line there is never any resource to spare. **(c)-(e)** Show control expenditure and disease progress for parameter sets corresponding to points A, B and C (as marked on panel **(a)**), illustrating the range of behaviour that is possible as the maximum budget is changed. In all cases, the switching time is marked with a dotted vertical line. Note that the switch in priority in Case C does not affect the control that is carried out, since the budget is not limiting at the time in question.

Epidemics spreading slowly are relatively easy to manage, and so can be controlled simply by always treating in the high value region. High infection rates give epidemics that spread rapidly, and so the more important high value region must again always be prioritised to keep the epidemic under control there. At intermediate spread rates, however, disease spreads slowly enough to allow reduction of infectious pressure by initially treating in the buffer region, but sufficiently quickly that such a reduction of pressure from the buffer is necessary to ensure an optimal result.

The diagonal “kink” in the shading in Figure 4a corresponds to a change in optimal control regime. At high maximum budgets the budget never limits the amount of control that can be carried out(above the diagonal kink). At very low maximum budgets control resources are always limited and the optimal control must distribute the resource appropriately. There is an intermediate region where the optimal control transforms between the two regimes.

#### Effect of the maximum rate of control

The effect of the maximum control rate *η* is shown in Figure 4b, revealing the value of this parameter also affects which values of the rate of infection(*β*) require a switch of focus in the optimal control strategy. As control becomes potentially more effective through faster maximum rates of treatment, the intermediate range over which a switching strategy is optimal shifts to faster spreading epidemics which have higher infection rates.

#### Effect of the initial level of infection and the external infectious pressure

Figure 5 shows the response of the switching time to the infection rate, *β*, the number of initially infected hosts in the buffer region, *I*_*B*_(0), and the external force of infection, 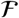 (recall in our model external inoculum can only infect the buffer region).

**Figure 5.**
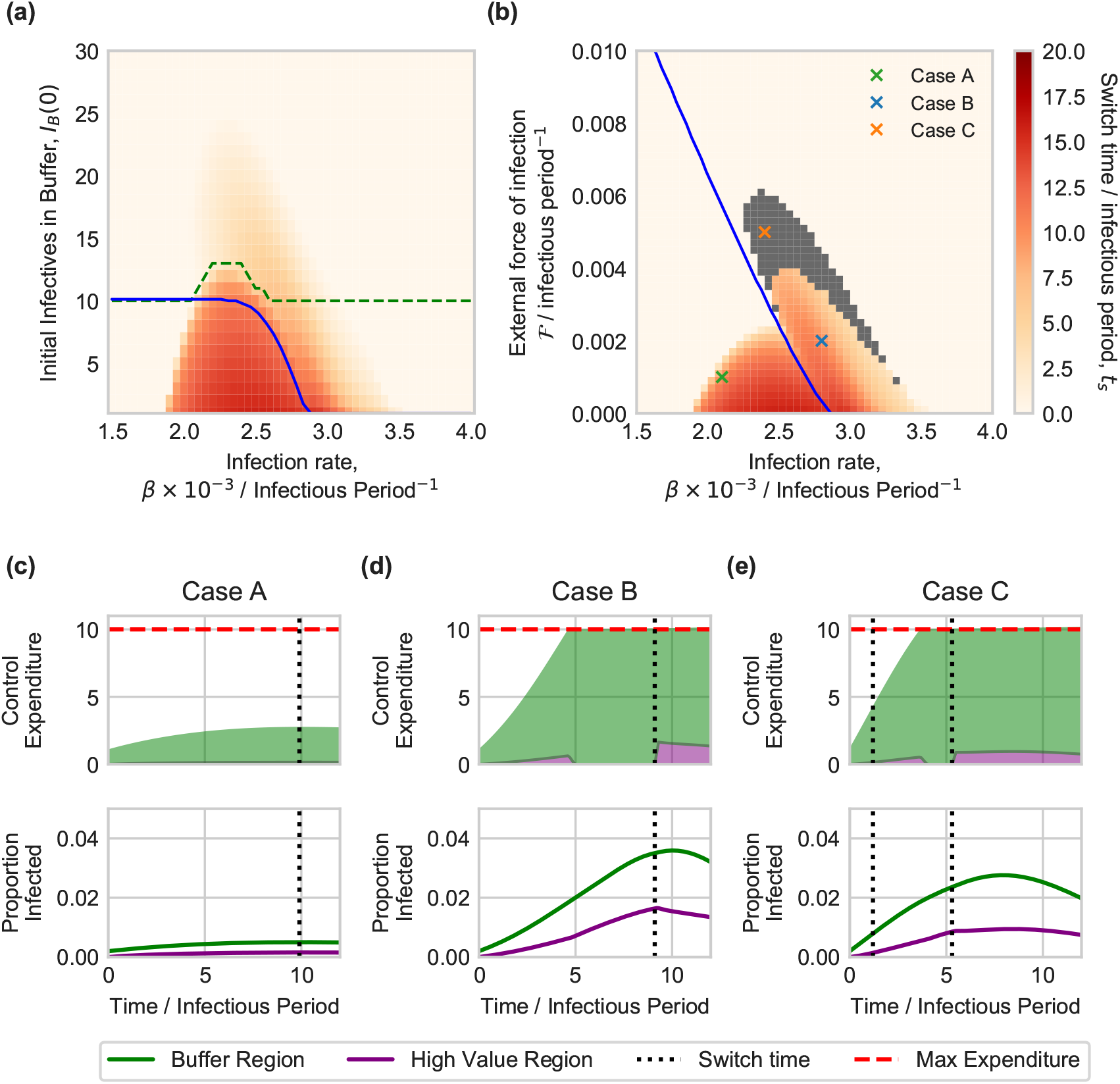
Effect of initial level of infection within, and force of infection upon, the buffer region. Parameter scans were run relative to a baseline set by the default parameters in Table 1, although using a longer timescale of interest(*T* = 20) to allow patterns in switching times to be more clearly seen. **(a)** shows the switch time as a function of the infection rate, *β*, and the initial number of infected hosts in the buffer region, *I*_*B*_(0). The blue line indicates where the budget becomes limiting on control resources, i.e. below the blue line maximal control can be carried out in both regions for the full epidemic. The green dashed line indicates when the optimal control switch occurs whilst the budget is limiting, i.e. above this line when priority switches to the value region, the budget is still limiting. **(b)** shows the effect of the infection rate, *β*, and the external force of infection on the buffer region, 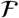 (with a single initially infected host in the buffer region, *I*_*B*_(0) = 1). The blue line has the same meaning as in(a). The grey region highlights of parameters for which multiple switches are identified. **(c)-(e)** Show control expenditure and disease progress for parameter sets corresponding to points A, B and C (as marked on panel **(a)**), illustrating the range of behaviour that is possible as the external force of infection is increased(see main text). In all cases, any switching time is marked with a dotted vertical line; note that for the single switch in Case A, and the first of the two switches in Case C, since the budget is not limiting at the time in question, the switch in priority does not affect the control that is carried out.

At low levels of initial infection in the buffer and/or external force of infection, the switching strategy is once again optimal at intermediate infection rates. However, as the rate of invasion of the buffer is increased, through either more initial infection or via a higher external force of infection, the range of infection rates over which a switching strategy is optimal decreases. For sufficiently high rates of invasion in the buffer region(i.e. moving up the *y* axis in Figures 5a,b) a switching strategy becomes sub-optimal, because the disease spreads faster in the buffer, and so control is less effective there than in the high value region.

The particular shapes of the responses in Figures 5a,b are affected by where the maximum expenditure rate becomes limiting, with the budget becoming more constraining towards the upper right corners. Whilst we do not attempt to explain the underlying drivers for the optimal control allocation for each parameterisation, the pattern of optimal control does change when the budget no longer allows maximal control for the whole epidemic. Note that for some values of the external force of infection, optimal strategies with additional switches are found; these are the grey regions in Figure 5b. As described already, these switches occur whilst the budget is not limiting, and so have no effect on the control realised.

To illustrate the full range of possibilities for the optimal control strategy, we focus on three particular cases taken from Figure 5b. Case A finds a switch but as the budget is not limiting, it has no effect. In case B the budget is limiting so the switch has an effect. In case C there is an additional switch early in the epidemic, but since the budget is not limiting at that point it has no effect.

### Optimal strategies can be robust to model misspecification, but requires some prior knowledge of transmission dynamics

Finally, we test how these optimal strategies perform when the parameters controlling the underlying model are not known accurately(Figure 6). We do this by introducing a systematic error into the infection rate of the model used to optimise the control strategy, varying the value of *β* from 50% smaller to 50% larger than the “true” value. The resulting optimal control specifies an expenditure over time (*f*_*i*_(*t*)*I*_*i*_(*t*) for each region) which is applied to a model with the “true” infection rate. Large errors in the infection rate lead to worse control of the epidemic. It is also important to note that for large underestimates of the infection rate, the optimised control is worse than simply allocating the full budget to the high value region, with no treatment in the buffer region. The optimised control strategies lead to wasted resources that could be better allocated to the other region(Figure 6(b,c)). These results suggest that if prior knowledge of parameters controlling transmission is imprecise, simpler control strategies might be better(***Hyatt-Twynam et al., 2017***).

**Figure 6.**
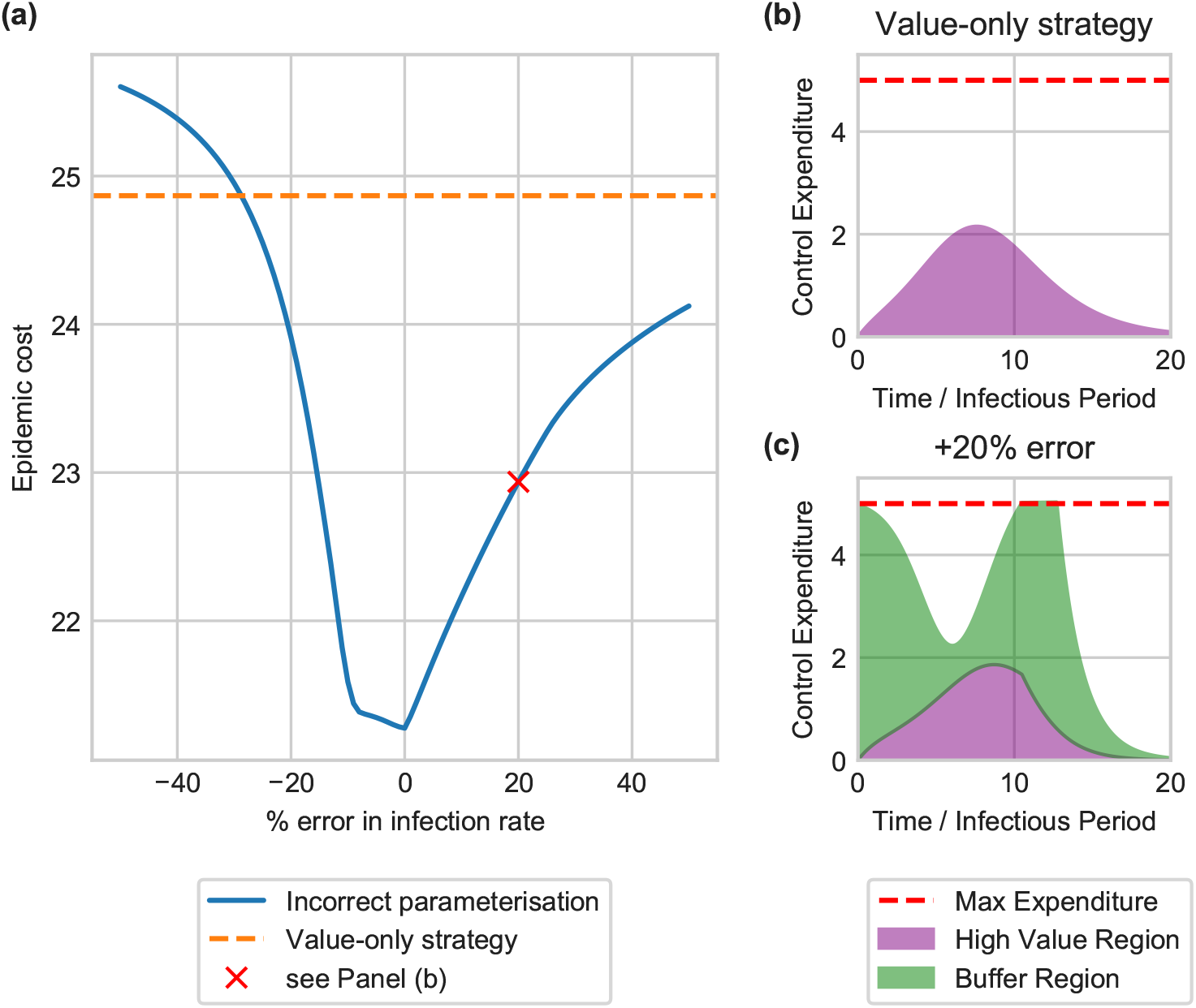
Effect of incorrect parameterisation on the performance of optimal control strategies. Control is optimised using an incorrect infection rate, and the budget allocations applied to the correct model. The baseline parameters use the default values, with an infection rate of 0.0028 host^−1^ t^−1^ and a maximum expenditure rate of 5 hosts, the same values as used for the switching strategy in Figure 3. **(a)** shows the epidemic cost as the percentage error in the infection rate is varied. Larger errors lead to worse performance, and can be worse than the simple strategy of full allocation to the high value region(shown in **(b)). (c)** shows the wasted control resources using an infection rate overestimated by 20 %. Here too much control is allocated to the high value region that cannot be spent, and so is wasted. Similarly, not enough resources are allocated when the infection rate is underestimated, also leading to wasted control resources.

## Discussion

Landscape scale control of sudden oak death in California has not been possible for some years(***Cunniffe et al., 2016***). However highly valuable sub-populations at risk due to the epidemic – such as areas important for tourism or cultural reasons, or for ecological reasons to conserve biodiversity – might yet be protected. Slowing spread to these regions is itself valuable(***Cobb et al., 2013***). An obvious question is then how this goal can best be achieved. This requires partitioning limited resources for disease management between the population of particular interest and the growing epidemic elsewhere, and determining whether this partitioning should vary with time and with the current state of the epidemic(***Rowthorn et al., 2009***). This must be done in a situation in which different sets of stakeholders have different objectives(***Craig et al., 2018***), but must nevertheless work together(***Laranjeira et al., 2020***).

We used the outbreak of sudden oak death in Humboldt County, California to motivate our work. After developing a simple mathematical model representing disease within the “high value” region of the Redwood National Park and in a “buffer” region surrounding it, we used optimal control theory to find the most effective time-varying allocation of a limited budget for disease management between these two regions. To minimise the final amount of infection in the National Park, we found it is very often better to prioritise exclusively that area for treatment(Figure 2). However, while it is clearly very intuitive, we have identified that this strategy is not always optimal. It can instead be better to start by prioritising disease control in the buffer region, and only to switch to prioritising the high value region later in the epidemic(Figure 3). We have found that such a “switching” strategy is most likely to be optimal for intermediate values of the infection rate(Figure 3).

This type of policy involving switching attention from reducing a long term threat to focusing on short term gains has been identified before. For example, ***Hastings et al. (2006)*** used a class-structured model to investigate control of *Spartina alterniflora*(an invasive species of deciduous grass) in Willapa Bay, Washington. The key prediction was that early control within each season should preferentially remove plants from the class with the highest reproductive value, only shifting later to remove plants that contribute most to the next season’s population. Similar switching strategies have also been found in other studies relating to management of invasive species(***Carrasco et al., 2009***).

Switching strategies have also been identified in other epidemiological models. For example, ***Ndeffo Mbah and Gilligan (2011)*** found that disease management across sub-populations is most efficient when resources are first allocated to whichever group is more infected, and later switching to treat the less infected group. A similar strategy was even found by ***Ndeffo Mbah and Gilligan (2010b)*** for management of sudden oak death. Switching strategies have also been found for distribution of prophylactic vaccines in a spatially structured population(***Keeling and Shattock, 2012***), and switching between immunisation and palliative care during an epidemic when resources are limited(***Klepac et al., 2012***).

For the simple model we considered here, the optimal strategies could in fact probably have been identified by an exhaustive scan over switching times. However for more complex models with more complex switching strategies this will not generally be true. Whilst the setting was highly simplified, we can already begin to see some of the potential limitations of these more complex strategies for practical use. Although not included in our analysis, there are additional costs associated with changing policy during an epidemic, i.e. switching region priorities, and these costs must be balanced with the benefit to epidemic control. These costs could be included in the objective function as optimised, for example by adding a term penalising rapidly changing controls(***Clarke et al., 2013***).

Direct use of optimal control theory also facilitated the extensive scans over alternative parameterisations of our model(Figures 3 and 5), allowing our relatively intuitive characterisation of a number of outcomes based on which region is prioritised. Although some work in optimal control of plant disease does translate strategies for practical application, and consider the impact of parameters(e.g.(***Hamelin et al., 2021***)), more often results are presented for a single parameterisation and tend not to focus on the practical implementability of the strategies that result(***Bokil et al., 2019***; ***Basir et al., 2017***; ***Basir and Ray, 2020***; ***Chen-Charpentier and Jackson, 2020***; ***Alemneh et al., 2021***).

We have also shown how imprecision in knowledge of parameters controlling disease spread can lead to less effective disease management(Figure 6). Indeed in such cases it may even be better to use the simplest possible strategy rather than use optimal control theory at all. Consideration of parameter uncertainty when determining the optimal strategy is important(***Epanchin-Niell and Hastings, 2010b***). The type of control strategy identified here, a bang-bang switching control, can be particularly sensitive to precise knowledge about the optimal switch time. ***Forster and Gilligan***(***2007***) showed that when applying control optimised using a mean-field model of an epidemic to a spatially-explicit model, errors in the switch time can lead to performance that is worse than a simple constant strategy. A study by ***Carrasco et al.***(***2009***) showed that when controlling invasive species, the optimal strategy when parameters are known precisely is not always optimal when parameter uncertainty is introduced. This also echoes results from so-called “risk-based” control strategies, which attempt to use additional epidemiological information to develop precise control strategies. When knowledge is limited simpler control strategies are very likely to be more appropriate(***Hyatt-Twynam et al., 2017***)

While here we have focused on the particular example of sudden oak death, we note our model is sufficiently abstract that, at least in principle, it could apply in other settings. Control by artificially shortening hosts’ infectious period could correspond to, for example, antibiotic treatment of agricultural animals, or encouraging individuals to self-isolate following symptoms of human respiratory diseases. Analogous problems in which a particular region or sub-population must be protected in the face of a ongoing epidemic also arise rather naturally for other pathosystems. For example, for COVID-19 in Western Europe, an obvious focus before the effects of vaccination became apparent lay in reducing numbers of infections of elderly residents of care facilities, since older individuals tend to suffer much worse outcomes following infection(***Palmer et al., 2021***).

We concentrated here upon only the relative amounts of one single control – i.e. removing infected hosts – in each of our two regions. For sudden oak death, a number of other management strategies are sometimes recommended(***Swiecki and Bernhardt, 2013***), and these were not included in our model. In particular, we did not consider pre-emptive removal of “spreader” host species such as California bay laurel that drive local epidemics. Such “thinning” is well-acknowledged to reduce the risk of *P. ramorum* infection entering an individual stand, or the rate at which it spreads following invasion(***Bussell and Cunniffe, 2020***). Our main motivation to only consider a single control strategy was simplicity: two distinct controls(i.e. reactive removal of infected hosts and pre-emptive removal of healthy hosts) would immediately double the complexity of the optimization problem(*cf.* Equation 2). It would also increase, perhaps significantly, the complexity of the optimal strategies that would need to be interpreted. We also note that preemptive removal of healthy hosts is somewhat specific to plant diseases in natural environments and so including it here would dilute the generic nature of our study. Finally, species-specific host removal rates would be difficult to represent in the type of model used here, in which all potentially infected species are amalgamated into a single category of “hosts”(***Meentemeyer et al., 2011***; ***Thompson et al., 2016a***). However, we note preemptive removal of certain host species has been included in more detailed system-specific models of sudden oak death in particular, including our own(***Bussell and Cunniffe, 2020***), and(perhaps unsurprisingly) suggest the focus of control should switch from preemptive to reactive treatments as any local epidemic becomes larger.

This paper is intended to be an illustrative analysis, highlighting the epidemiological principles at play when protecting a valuable sub-population in the face of an otherwise unmitigated epidemic. We therefore deliberately simplified the wide range of management strategies available for sudden oak death, and instead concentrated in detail on a single type of management, i.e. removing infected hosts. However this was sufficient for us to show that the obvious benefit of direct treatment of the most valuable region might be outweighed by the delayed benefit from treating elsewhere(***Rowthorn et al., 2009***). However, as illustrated by our analysis, the optimal strategy depends on the complex interplay between the epidemiological parameters, the level of precision with which these parameters are known, the budget available for control and the current state of the epidemic. For practical application, our model would need to be extended to more faithfully capture sudden oak death epidemiology, for which cryptic infection, long-distance dispersal, epidemiological differences between different host species, spatial structure, alternate control strategies and stochasticity are all likely to be important. Given that, at least under some circumstances, the cost of protecting the high-value region is felt by stakeholders elsewhere, accounting for stakeholder behaviour could also be an interesting extension to the modelling work presented here(***Milne et al., 2020***; ***Murray-Watson et al., 2021***). However recent work has shown how insights derived via optimal control theory could nevertheless potentially be applied irrespective of any or all of these complexities(***Bussell et al., 2019***; ***Bussell and Cunniffe, 2020***). By showing how non-intuitive control strategies might arise from such an exercise, the key contribution of this paper is to develop a framework for identifying broad categories of control strategy to further investigate in such a setting.

## Author Contributions

E.H.B. and N.J.C. designed the study, E.H.B. conducted the analysis and wrote the initial draft of the manuscript. Both authors contributed to data interpretation, manuscript editing and discussion.

## Acknowledgements

We thank Jane White and Olivier Restif, the examiners of the PhD thesis upon which this paper is based, for useful discussions of the work.

## Data Accessibility

A reference implementation of our underlying model and optimisation procedure is available at https://github.com/ehbussell/PatchOptimalControl.

