## Supplementary Material S1 Text for "Optimal strategies to protect a sub-population at risk due to an established epidemic"

### Supporting Information S1 Text

#### S1 Effect of a growing epidemic in the generally infested region

Our model assumes a constant external force of infection on the buffer region (the parameter $\mathcal{F}$ in Equation 1 of the main text). This can be generalised to instead represent a growing epidemic in the generally infested region via an exponentially increasing form

$$\mathcal{F}' = F_0 \exp(rt), \qquad \text{(App.1)}$$

in which $F_0$ corresponds to the initial size of the external epidemic.

In Figure 1 epidemic dynamics are shown for the default parameters using the standard constant external force of infection (solid lines) and an exponentially increasing external force of infection (dashed lines). Under no control the change has very little effect on the overall dynamics since infection rates are dominated by within-region infection rates. The growth rate is taken to be $r = 0.1$ t$^{-1}$, with the initial value $F_0$ set to ensure that the integrated force of infection over time is the same as for the constant force of infection, i.e.

$$F_0 = \frac{r\mathcal{F}T}{e^{rT} - 1}, \qquad \text{(App.2)}$$

in which $T$ is the simulation time.

Under control prioritising the high value region, and using a smaller infection rate of $\beta$ =0.003 host$^{-1}$ t$^{-1}$, there is a small difference in epidemic dynamics (Figure 2). The overall pattern of dynamics is very similar though.

We conclude that slowly growing epidemics in the generally infested region, or at least growth that has little impact on the force of infection in the buffer region, is likely to have little impact on our results. However, the assumptions in the patch-based model we use in the main paper will break down if the generally infested region is very close to the buffer region, or if the force of infection changes significantly over the timescale of the epidemic in the buffer region.

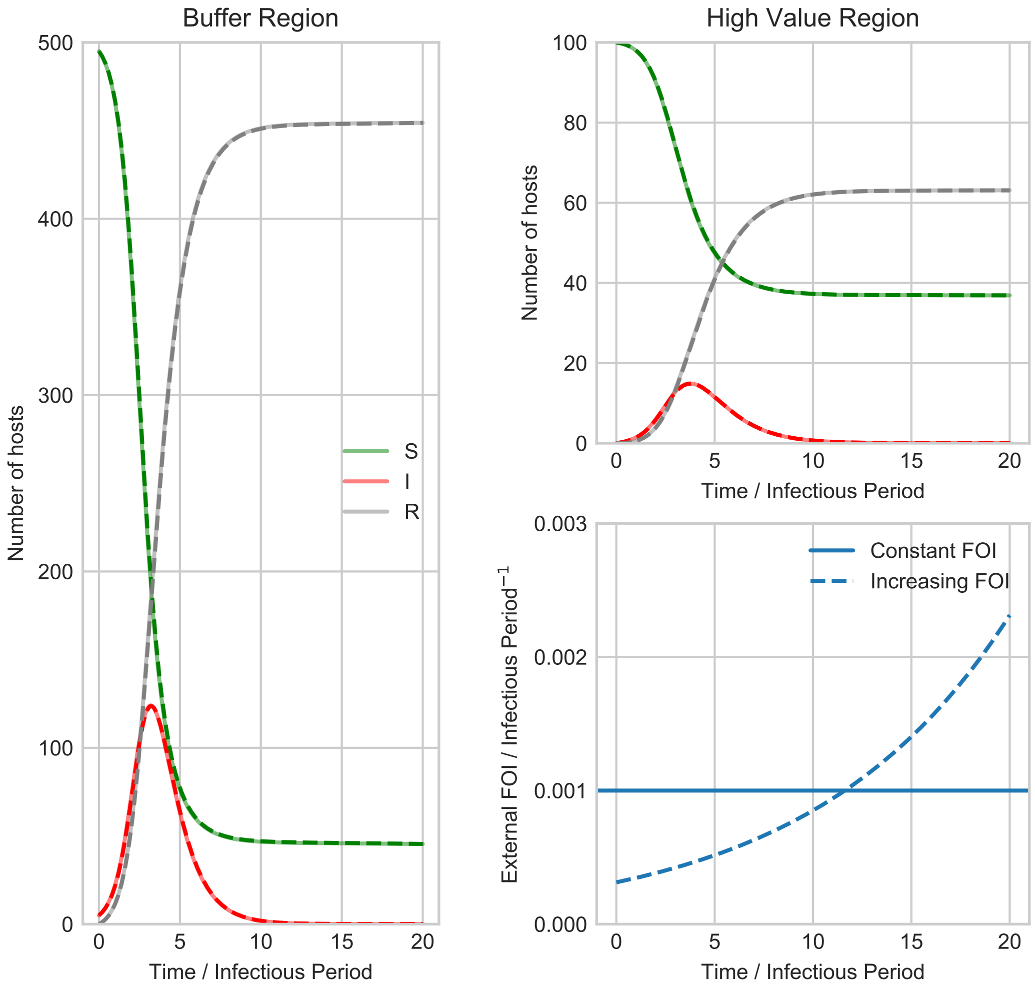


**Figure 1. A growing epidemic in the generally infested region** Using the default parameters, there is no visible difference between the model dynamics with a constant external force of infection (solid lines; $\mathcal{F}$ is constant as used in the main paper), and an exponentially growing force of infection (dotted lines; $\mathcal{F}' = F_0 \exp(rt)$, where $r = 0.1$ per infectious period, and where $F_0$ is normalised according to Equation App.2).

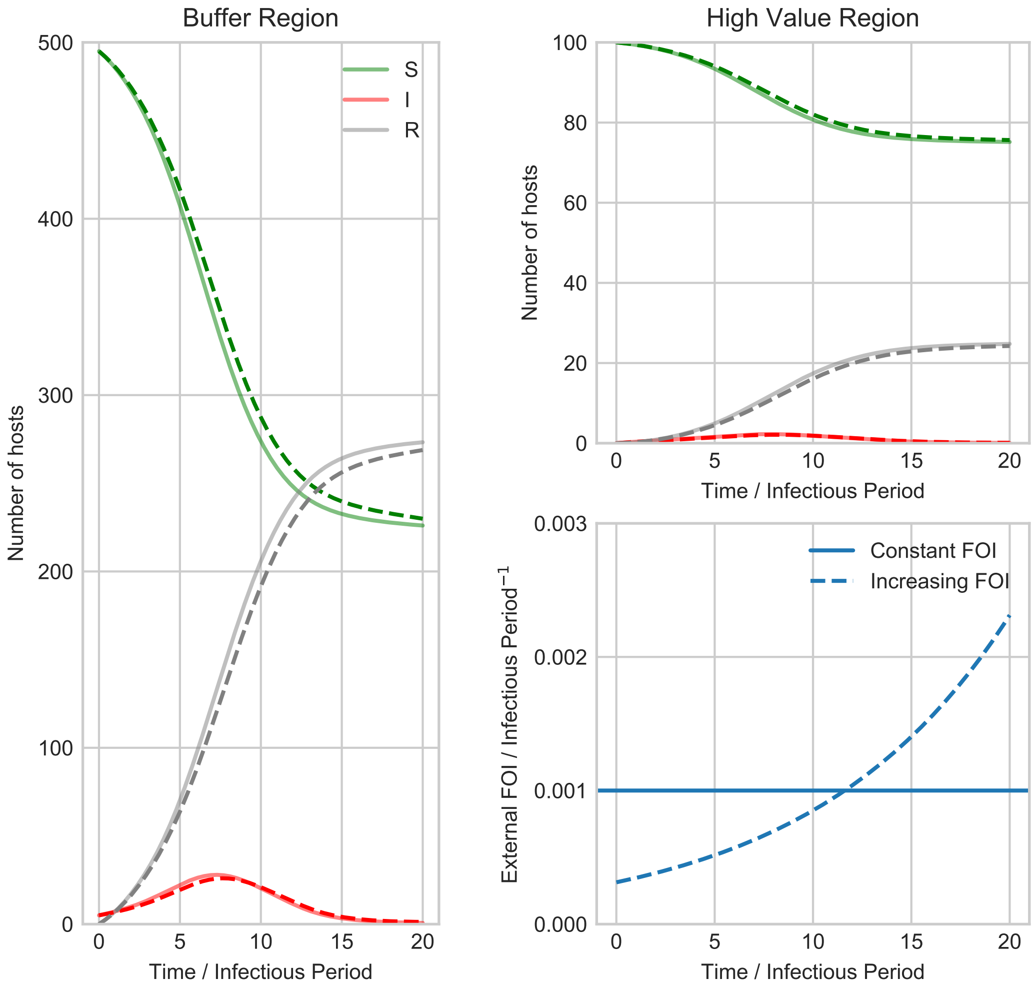


**Figure 2. When the external epidemic is more significant, the approximation is slightly less precise**. We consider a case in which there is a smaller within-region infection rate ($\beta$ =0.003 host$^{-1}$ t$^{-1}$ rather than $\beta$ =0.005 host$^{-1}$ t$^{-1}$) and when control is being done in the high value region. Since the dynamics of the external forcing are then more important in relative terms, there are visible differences between the results of the two models. However the differences in fact remain relatively small over the full time course of the epidemic.
